# Temporal signatures of criticality in human cortical excitability as probed by early somatosensory responses

**DOI:** 10.1101/809285

**Authors:** T. Stephani, G. Waterstraat, S. Haufe, G. Curio, A. Villringer, V. V. Nikulin

## Abstract

Brain responses vary considerably from moment to moment, even to identical sensory stimuli. This has been attributed to changes in instantaneous neuronal states determining the system’s excitability. Yet the spatio-temporal organization of these dynamics remains poorly understood. Here we test whether variability in stimulus-evoked activity can be interpreted within the framework of criticality, which postulates dynamics of neural systems to be tuned towards the phase transition between stability and instability as is reflected in scale-free fluctuations in spontaneous neural activity. Using a novel non-invasive approach in 33 male participants, we tracked instantaneous cortical excitability by inferring the magnitude of excitatory post-synaptic currents from the N20 component of the somatosensory evoked potential. Fluctuations of cortical excitability demonstrated long-range temporal dependencies decaying according to a power law across trials – a hallmark of systems at critical states. As these dynamics covaried with changes in pre-stimulus oscillatory activity in the alpha band (8–13 Hz), we establish a mechanistic link between ongoing and evoked activity through cortical excitability and argue that the co-emergence of common temporal power laws may indeed originate from neural networks poised close to a critical state. In contrast, no signatures of criticality were found in subcortical or peripheral nerve activity. Thus, criticality may represent a parsimonious organizing principle of variability in stimulus-related brain processes on a cortical level, possibly reflecting a delicate equilibrium between robustness and flexibility of neural responses to external stimuli.

**Significance Statement:** Variability of neural responses in primary sensory areas is puzzling, as it is detrimental to the exact mapping between stimulus features and neural activity. However, such variability can be beneficial for information processing in neural networks if it is of a specific nature, namely if dynamics are poised at a so-called critical state characterized by a scale-free spatio-temporal structure. Here, we demonstrate the existence of a link between signatures of criticality in ongoing and evoked activity through cortical excitability, which fills the long-standing gap between two major directions of research on neural variability: The impact of instantaneous brain states on stimulus processing on the one hand and the scale-free organization of spatio-temporal network dynamics of spontaneous activity on the other.

## 1 Introduction

Neural responses are characterized by remarkable variability, even to identical physical stimuli. This well-known phenomenon has been attributed to fluctuations of the neuronal network’s state (Arieli et al., 1996; Sadaghiani et al., 2010), observable via diverse neuronal measures such as EEG (Romei et al., 2008; Vanrullen et al., 2011; Rahn and Basar, 1993; Iemi et al., 2019; Forschack et al., 2017), BOLD signal (Fox and Raichle, 2007; Becker et al., 2011), local field potentials (Arieli et al., 1996), and single-cell recordings (Azouz and Gray, 1999; Churchland et al., 2010). So far, studies on neuronal variability have mainly focused on the strength of variability (Dinstein et al., 2015; Garrett et al., 2013; Churchland et al., 2010). The dynamics of network states over time, however, have often been neglected despite their ability to provide further insights into the underlying spatio-temporal organization principles.

In this context, a certain type of fluctuation pattern known as *power-law dynamics*, which indicates that a signal possesses scale-free properties, is of particular interest. Such power-law relationships represent a hallmark of the (self-)organization of complex systems at a *critical state* (Sethna et al., 2001; Muñoz, 2018), the point of a phase transition between two distinct system regimes, such as order and disorder (Beggs and Plenz, 2003; Bak et al., 1987; Bak et al., 1988), at which the dynamic range, information processing and capacity of a system are maximized (Kinouchi and Copelli, 2006; Shew and Plenz, 2013). Figure 1A visualizes these system configurations using the *Ising model of ferromagnetism* (Ising, 1925).

**Figure 1.**
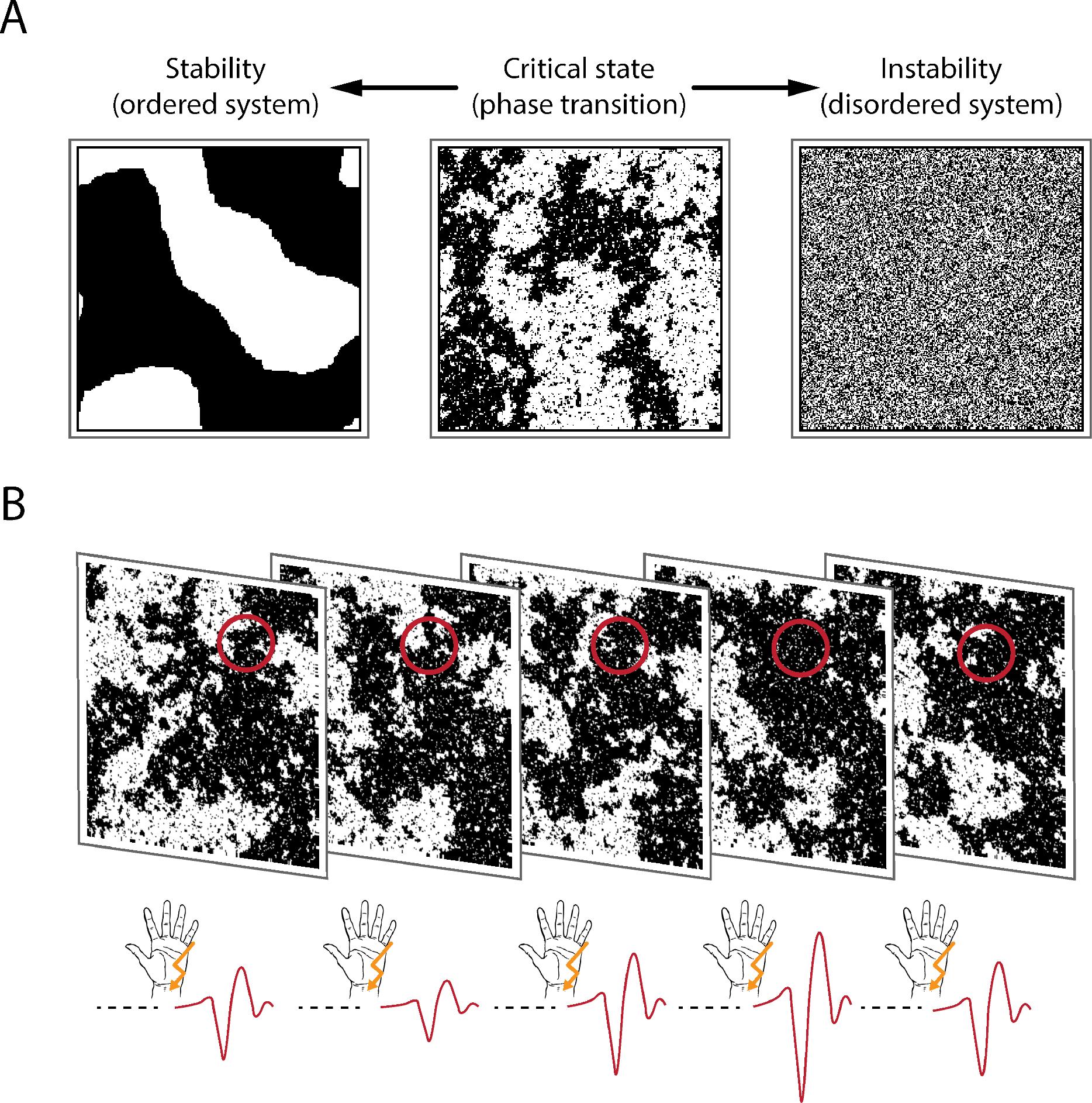
Types of system dynamics and how to probe them. A) *Ising model* at different system states: ordered, critical, disordered (from left to right). Transferred to a grid of neurons, black and white shades reflect firing and non-firing neurons, or, as in the context of our study, neurons that can or cannot be recruited by the stimulus. Here, snapshots of the system at a given point in time are shown. In an ordered system, local interactions dominate and lead to highly stable neural activity. In contrast, firing patterns in a disordered system are highly unstable and quickly change from moment to moment in a stochastically independent manner (i.e., white noise). At the critical state, the system resides at the border between the tendencies either towards an ordered or towards a disordered system. This is reflected by the spatio-temporal dynamics, that is, scale-invariance or power-law dynamics. Scale-invariance is visible from the middle panel since similar clusters of black pixels occur on all scales. B) Experimental paradigm. The instantaneous state of the neuronal system (illustrated here with snapshots of the Ising model at the critical state) is probed by somatosensory stimuli. The amplitude of the N20 component of the SEP is expected to be proportional to the number of neurons that can be recruited by the stimulus at a given moment (black pixels in the probed area, which is marked by the red circles), therefore reflecting a measure of instantaneous cortical excitability. In a stable system, the number of neurons that can be excited would barely change over time, whereas in an unstable system this would vary randomly. However, at the critical state configurations would show the largest range of variation over time with fluctuations following a temporal power law.

Empirically, power-law dynamics have been found in the size and duration of neuronal avalanches of various species, such as rats (Beggs and Plenz, 2003; Friedman et al., 2012), monkeys (Petermann et al., 2009; Yu et al., 2017), zebrafish larvae (Ponce-Alvarez et al., 2018), and humans (Priesemann et al., 2013; Shriki et al., 2013; Arviv et al., 2015). In the temporal domain, power-law dynamics can be observed in human resting state fMRI networks (Tagliazucchi et al., 2013), as well as in amplitude fluctuations of alpha and beta band activity by way of MEG/EEG (Linkenkaer-Hansen et al., 2001; Palva et al., 2013), associated with long-range temporal dependencies. Thus, criticality may represent a general and appealingly parsimonious explanation of neuronal variability in the brain. However, the functional link between critical dynamics, fluctuations of ongoing neural activity, and cortical excitability remains elusive – particularly for the human brain.

To establish this link, we developed an approach to probe instantaneous cortical excitability on a single-trial level using somatosensory evoked potentials (SEP) in EEGs on humans in response to electrical median nerve stimuli (Fig. 1B). Specifically, the N20 component of the SEP is thought to solely reflect excitatory post-synaptic potentials (EPSPs) of the first thalamo-cortical volley (Wikström et al., 1996; Nicholson Peterson et al., 1995; Bruyns-Haylett et al., 2017), generated in the anterior wall of the postcentral gyrus, Brodmann area 3b (Allison et al., 1991). Therefore, the amplitude of this early part of the SEP represents a direct measure of the instantaneous excitability of a well-defined neuronal population in the primary somatosensory cortex. Additionally, to bridge the gap between evoked and ongoing neuronal activity, we evaluated pre-stimulus oscillations in the alpha band (8–13 Hz) of the same neuronal sources, a classical index of cortical excitability in ongoing neural activity (Klimesch et al., 2007; Romei et al., 2008).

The temporal structure of these two measures of excitability demonstrated power-law dynamics over time, which were entirely generated on the cortical level as shown by control analyses of subcortical and peripheral signal variability. Furthermore, the SEP-derived measure of cortical excitability and pre-stimulus alpha band activity were coupled regarding both their amplitudes and power-law exponents. Therefore, variability in both ongoing and evoked neural activity is likely to be governed by the same near-critical network dynamics.

## 2 Materials and Methods

### Participants

EEG data were recorded from 33 male human subjects. Two subjects were excluded because no clear SEPs were visible in the single-trial analysis, probably due to suboptimal placement of the stimulation electrodes and a low SNR of the EEG. The remaining sample of 31 subjects had an average age of *M* = 26.9 years (*SD* = 5.0). All participants were right-handed (lateralization score, *M* = +92.9, *SD* = 11.7), as assessed using the Edinburgh Handedness Inventory (Oldfield, 1971), and did not report any neurological or psychiatric disease. All participants gave informed consent and were reimbursed monetarily. The study was approved by the local ethics committee.

### Stimuli

Somatosensory stimuli were applied using electrical stimulation of the median nerve. A non-invasive bipolar stimulation electrode was positioned on the left wrist (cathode proximal). The electrical stimuli were designed as squared pulses of a 20- μs duration. The stimulus intensity was set to 1.2 x motor threshold, leading to clearly visible thumb twitches for every stimulus, as individually determined by a staircase procedure prior to the experiment. Stimuli were applied using a DS-7 constant-current stimulator (Digitimer, Hertfordshire, United Kingdom).

### Procedure

During the experiment, participants were seated comfortably in a chair their hands extended in front of them in the supinate position on a pillow. Electrical stimuli were presented in a continuous sequence with inter-stimulus intervals (ISI) ranging from 713 to 813 ms (randomly drawn from a uniform distribution; ISI_average_ = 763 ms). In total, 1000 stimuli were applied, divided into two blocks of 500 stimuli each with a short break in between. Participants were instructed to relax and fixate their gaze on a cross on a computer screen in front of them while receiving the stimuli.

### Data Acquisition

EEG data were recorded from 60 Ag/AgCl electrodes at a sampling rate of 5000 Hz using an 80-channel EEG system (NeurOne, Bittium, Oulu, Finland) with a bandwidth of 0.16 to 1250 Hz. Electrodes were mounted in an elastic cap (EasyCap, Herrsching, Germany) at the international 10-10 system positions FP1, FPz, FP2, AF7, AF3, AFz, AF4, AF8, F7, F5, F3, F1, Fz, F2, F4, F6, F8, FT9, FT7, FT8, FT10, FC5, FC3, FC1, FC2, FC4, FC6, C5, C3, C1, Cz, C2, C4, C6, CP5, CP3, CP1, CPz, CP2, CP4, CP6, T7, T8, TP7, TP8, P7, P5, P3, P1, Pz, P2, P4, P6, P8, PO7, PO3, PO4, PO8, O1, and O2. Four additional electrodes were placed at the outer canthus and the infraorbital ridge of each eye to record the electro-oculogram (EOG). During recording, the EEG signal was referenced to FCz, and POz served as ground. All impedances were kept below 10 kΩ. For source reconstruction, EEG electrode positions were measured in 3D space individually for each subject using Polhemus Patriot (Polhemus, Colchester, Vermont). Additionally, the compound nerve action potential (CNAP) of the median nerve was measured using two bipolar electrodes, positioned at the inner side of the left upper arm.

Structural T1-weighted MRI scans (MPRAGE) of each participant were obtained during a different testing date prior to the experiment, on a 3T Siemens Verio, Siemens Skyra or Siemens Prisma scanner (Siemens, Erlangen, Germany).

### EEG pre-processing

Stimulation artifacts were cut out and interpolated between −2 to 4 ms relative to stimulus onset using Piecewise Cubic Hermite Interpolating Polynomials (PCHIP). The EEG data were band-pass filtered between 30 and 200 Hz, sliding a 4^th^ order Butterworth filter forwards and backwards over the data to prevent phase shift. With this filter, we specifically focused on the N20-P35 complex of the SEP. Furthermore, this filter effectively served as baseline correction of the SEP since it removed slow trends in the data, reaching an attenuation of 30 dB at 14 Hz, thus ensuring that fluctuations in the SEP did not arise from fluctuations with slower frequencies (e.g., alpha band activity). (The relationship between decibels [dB] and magnitude is defined as *dB* = 20 * *log*_10_[*magnitude*]). Bad segments of the data were removed by visual inspection, resulting in 989 trials on average per participant. The data were then re-referenced to an average reference. Eye artefacts were removed using independent component analysis (ICA). For analysis of SEPs, the data were segmented into epochs from −100 to 600 ms relative to stimulus onset. EEG pre-processing was performed using EEGLAB (Delorme and Makeig, 2004), and custom written scripts in MATLAB (The MathWorks Inc., Natick, Massachusetts).

### Single-trial extraction using CCA

Single-trial SEPs were extracted by applying a variant of Canonical Correlation Analysis (CCA), as previously proposed by Waterstraat et al.. CCA is used for finding weights *w_x_* and *w_y_* that mutually maximize the correlation between two signals *X* and *Y*, so that:

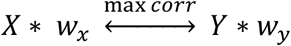

For extracting single-trial SEPs, we constructed *X* as a two-dimensional matrix (time by channel) containing all single-trial epochs (concatenated in the time domain), whereas *Y* contained the average SEP, concatenated as often as there were epochs (also concatenated in the time domain). The resulting weight matrix *w_x_* represents spatial filters that, in combination with *w_y_*, maximize the correlation between single-trial activity (*X*) and the average SEP (*Y*). To particularly focus on the early portion of the SEP, the spatial filters *w_x_* were trained using shorter segments from 5 to 80 ms post-stimulus but applied to the entire epochs from −100 to 600 ms. We derived a number of spatially distinct components by applying the spatial filters to the single-trial matrix, here denoted as *CCA components*:

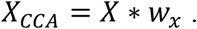

To characterize the CCA components in more detail, their spatial patterns were computed as

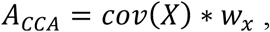

and components were visually identified that showed a tangential spatial pattern over the central sulcus as is typical for the N20-P35 complex (referred to as *tangential CCA components*). Furthermore, components were identified that showed a peak in the activity time course at 15 ms (referred to as *thalamic CCA components;* only in a subset of the sample). This procedure was performed individually for every subject for the first four CCA components, as sorted by their canonical correlation coefficients. Since CCA is insensitive to the polarity of the signal, the resulting tangential CCA components were standardized so that the N20 always appeared as a negative peak in the SEP (i.e., by inverting their spatial filters *w_x_*, if necessary). Furthermore, CCA is insensitive to the order of trials. Thus, the same spatial filters *w_x_* are obtained when permuting the order of single-trial SEPs and it is therefore not possible that CCA influences the temporal structure of SEP amplitudes across trials.

### SEP peak amplitudes and pre-stimulus oscillatory activity

N20 peak amplitudes were defined as the minimum value in single-trial SEPs of the tangential CCA components ±2 ms around the latency of the N20 in the within-subject average SEP. P35 peak amplitudes were defined accordingly as the maximum around the latency of the P35 in the within-subject average SEP.

Pre-stimulus alpha band activity was obtained from data segments between −200 to −10 ms relative to stimulus onset, band-pass filtered between 8 and 13 Hz (4^th^ order Butterworth filter forwards and backwards applied), after mirroring the pre-stimulus segments to both sides in order to reduce filter-related edge effects. To make a direct comparison with the SEP possible, we applied the spatial filter corresponding to the tangential CCA component to the pre-stimulus data. Subsequently, the pre-stimulus alpha envelope was measured by taking the absolute values of the signals processed with the Hilbert transform. To derive one pre-stimulus alpha metric for every trial, amplitudes of the alpha envelope were averaged across the whole pre-stimulus time window.

### EEG source reconstruction

To reconstruct the sources of the EEG signal, we estimated lead field matrices based on individual brain anatomies and individually measured electrode positions. The structural T1-weighted MRI images (MPRAGE) were segmented using the Freesurfer software (http://surfer.nmr.mgh.harvard.edu/), and a 3-shell boundary element model (BEM) was constructed which was used to compute the lead field matrix with OpenMEEG (Gramfort et al., 2010; Kybic et al., 2005). For two subjects, a template brain anatomy (ICBM152; Fonov et al., 2009) was used as no individual MRI scans were available. For one subject, standard electrode positions were used instead of individually measured positions. The lead field matrices were inverted using eLORETA (Pascual-Marqui, 2007), and sources were reconstructed for the spatial patterns of the tangential CCA component of every subject. Next, individual source spaces were transformed into a common source space based on the ICBM152 template using the spherical co-registration with the FSAverage atlas (Fischl et al., 1999) derived from Freesurfer, in order to average the obtained sources of the CCA components across subjects. The calculation of the individual head models and visualization of the sources was performed using Brainstorm (Tadel et al., 2011). The MATLAB implementation of the eLORETA algorithm was derived from the MEG/EEG Toolbox of Hamburg (METH).

### Processing of peripheral electrophysiological data (median nerve CNAP)

Analogously to the EEG data, stimulation artifacts were cut out and interpolated between −2 to 4 ms relative to stimulus-onset using Piecewise Cubic Hermite Interpolating Polynomials (PCHIP). Next, the data were high-pass filtered at 70 Hz, sliding a 4^th^ order Butterworth filter forwards and backwards over the data to prevent phase shift. Additionally, notch filters (4^th^ order Butterworth) were applied from 48 to 52 Hz and 148 to 152 Hz, respectively. epochs were extracted from −100 to 600 ms relative to stimulus onset.

### Detrended Fluctuation Analysis (DFA)

Power-law dynamics in the fluctuations of early SEPs as well as of pre-stimulus alpha band activity were quantified using Detrended Fluctuation Analysis (DFA; Kantelhardt et al., 2001; Hardstone et al., 2012). DFA calculates the fluctuation (i.e., standard deviation) of a cumulative signal on different time scales and tests whether its distribution follows a powerlaw: *F*(*τ*) ~ *τ^α^*, where *F* denotes the fluctuation function, *τ* the signal length (or window size), and *α* the power-law exponent. The DFA exponent *α* quantifies the extent of power-law dynamics of a signal, with *α* > 0.5 indicating persistent auto-correlations; whereas *α* = 0.5 is expected for a signal without a correlated temporal structure (i.e., white noise). We analyzed power-law dynamics in the fluctuation of SEP and pre-stimulus alpha amplitudes across trials with window sizes ranging from 7 to 70 trials, which correspond to time windows of 5.3 to 53.4 seconds. The same temporal window sizes were selected for the DFA of continuous alpha band activity.

### Evaluation of SNR

The signal-to-noise ratio of the single-trial SEP, as measured by the tangential CCA component, was quantified as the quotient of the root-mean-square signal in the time range of the SEP (10 to 50 ms) and a pre-stimulus baseline (−50 to −10 ms), so that 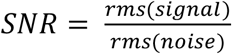.

The same procedure was applied to estimate the SNR of the CNAP and of the thalamic CCA component. For the CNAP we chose time windows from 5 to 8 ms and −8 to −5 ms, and for thalamic activity 12 to 18 ms and −18 to −12 ms, to estimate signal and noise, respectively.

### Simulation of the relationship between SNR and DFA exponent

Signals with DFA exponents systematically varying in the range from *α* = 0.5 to *α* = 0.8 were generated by filtering white noise with IIR filters whose coefficients depended on the desired DFA exponents as described in Schaworonkow et al. according to the algorithm of Kasdin. The length of these time series was set to 1000 data points corresponding to our empirical data from the SEP fluctuation across trials. These time series were mixed with white noise, that is, stochastically independent time series with DFA exponents of *α* = 0.5. The time series with varying DFA exponents were mixed with the noise at varying SNRs ranging from 0.001 to 6, defined as 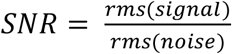. This procedure was repeated 100 times to account for the variance in the generation of random time series. Subsequently, DFA exponents of the mixed time series were measured and the average DFA exponent of the simulated signal was identified for which the SNR and DFA exponent of the mixed time series corresponded to our empirical analysis of SEP fluctuations.

### Simulation of the influence of temporal filtering on DFA exponents

To confirm that our temporal filtering did not cause the DFA exponent increases in the early SEP, we applied the same filtering to surrogate data with stochastically independent SEP fluctuations. SEP fluctuations across trials were simulated by decreasing or increasing an average SEP time course by a randomly generated factor for every trial. These signals were superimposed on continuous pink noise which was band-pass filtered between 30 and 200 Hz (4^th^ order Butterworth filter applied forwards and backwards), using a signal-to-noise ratio of 2, a typical value for empirical data. Subsequently, DFA was applied across trials for every time sample of the simulated SEP, corresponding to above described DFA analyses of the empirical SEPs.

### Statistical analyses

We compared the empirical DFA exponent time courses to surrogate data and applied cluster-based permutation tests to assess whether, and at which latencies, DFA exponents were significantly higher than it would be expected for stochastically independent fluctuation (i.e., white noise). First, we determined the expected DFA exponents for stochastically independent fluctuation by shuffling the trial order of our data and applying DFA to it. To account for variability due to random shuffling, this step was repeated 1000 times, and DFA exponents of these iterations were averaged, yielding an average surrogate DFA exponent time course for every subject. (Averaged across all samples and subjects, the mean DFA exponent was *α* = .512, thus slightly increased as compared to the theoretical DFA exponent of white noise of *α* = .5. This small empirical deviation may have been caused by the asymptotic behavior of DFA for small window sizes.) Next, the DFA exponents of the data with intact trial order were compared to the average DFA exponents of the surrogate data, using a two-sample t-test, resulting in a *t* value for every comparison over the time course of the SEP. To obtain clusters of increased DFA exponents, *t* values were thresholded at *p_pre_* = .001. Within clusters, *t* values were summed up to cluster *t* values *t_cluster,empirical_*. The same procedure was repeated 1000 times for the surrogate data, always comparing one surrogate dataset to the average surrogate data, which provided us with the distribution of cluster *t* values under the null hypothesis. Next, a cut-off value *t_cluster,crit_* was defined at the 99.9th percentile corresponding to a cluster threshold of *p_cluster_* =.001. Finally, *t_cluster,empirical_* of all clusters in the empirical DFA exponent time course were compared to *t_cluster,crit_* to identify clusters of significantly increased DFA exponents. With this procedure, we controlled for the number of samples over the SEP time course, inter-subject variability, and the distribution of amplitude values of the SEP from which DFA exponents were derived.

Analogously, DFA exponents of pre-stimulus alpha band activity were statistically tested using a t-test on group-level comparing them to the average DFA exponents of the null distribution, which was calculated from 1000 surrogate datasets with shuffled trial order. Similarly, the statistical significance of the DFA exponents of continuous alpha band activity was tested, however shuffling samples instead of trials to obtain DFA exponents under the null hypothesis.

To test the relationship of SNR and DFA exponents, we correlated the average SNR of single-trial SEPs with the area-under-the-curve (AUC) of DFA exponents between 10 and 50 ms post-stimulus, across participants using Spearman correlation.

Furthermore, we assessed the relationship between single-trial N20 peak amplitudes and pre-stimulus alpha amplitudes using a linear-mixed-effects model with *subject* as random factor, estimating the fixed effect as well as the random slope of the predictor *pre-stimulus alpha amplitude* with the dependent variable *N20 peak amplitude* (intercepts were included both for the fixed and random effects). Additionally, the relationship between N20 peak amplitudes and pre-stimulus alpha amplitudes was assessed with a permutation-based approach in which we compared their Spearman correlation coefficients with those from surrogate pre-stimulus alpha amplitudes with the same auto-correlated structure but shuffled phases generated by Adjusted Amplitude Fourier Transform (Theiler et al., 1992), as suggested in Schaworonkow et al.. Empirical correlation coefficients were averaged across subjects after Fisher’s Z transformation and compared with the null distribution of 10000 averaged correlation coefficients from the surrogate analyses to obtain the corresponding *p* value.

The covariation of N20 and P35 peak amplitudes was tested using a random-slope linear-mixed-effects model with *P35 peak amplitude* as dependent variable, *N20 peak amplitude* as independent variable and *subject* as random factor.

Finally, we correlated DFA exponents of pre-stimulus alpha activity and DFA exponents of the SEP. To account for the variability in the DFA exponent time course in the early SEP, we calculated the root-mean-square of DFA exponents in four subsequent time windows, 20 to 25, 25 to 30, 30 to 35, and 35 to 40 ms, and computed their Spearman correlation coefficients with the DFA exponents of pre-stimulus alpha activity, respectively. To control for the resulting multiple testing, we applied Bonferroni correction.

For all statistical analyses the significance level was set to *p* = .05. Correlation analyses as well as permutation-based statistics were performed in MATLAB (version 2017b, The MathWorks Inc., Natick, Massachusetts). The linear-mixed-effects models were calculated in R (version 3.5.1, R Core Team, 2018) using the lme4 (Bates et al., 2015) and denominator degrees of freedom were adjusted using Satterthwaite’s method (Satterthwaite, 1946) to derive a *p* value for the fixed effects as implemented in the R package lmerTest (Kuznetsova et al., 2017).

### Data and code availability

The data that supports the findings of this study are available upon request from the corresponding author (T.S.). The data cannot be made publicly available due to the privacy policies for human biometric data according to the European General Data Protection Regulation (GDPR).

The custom-written code that was used for data processing and statistical analyses is publicly available at https://osf.io/jzqdt/?view_only=dcb94617841445859adc5496d33c3cee.

## 3 Results

### SEPs and neuronal generators

The SEPs, averaged across all participants and trials, are shown in Figure 2A. The N20 component is visible as a negative peak at around 20 ms at electrodes contralateral to the stimulation site and posterior to the central sulcus. Furthermore, the scalp topography at 20 ms shows a tangential dipole centered over the central sulcus (Fig. 2B), consistent with the assumption of neuronal generators located in the anterior wall of the postcentral gyrus.

**Figure 2.**
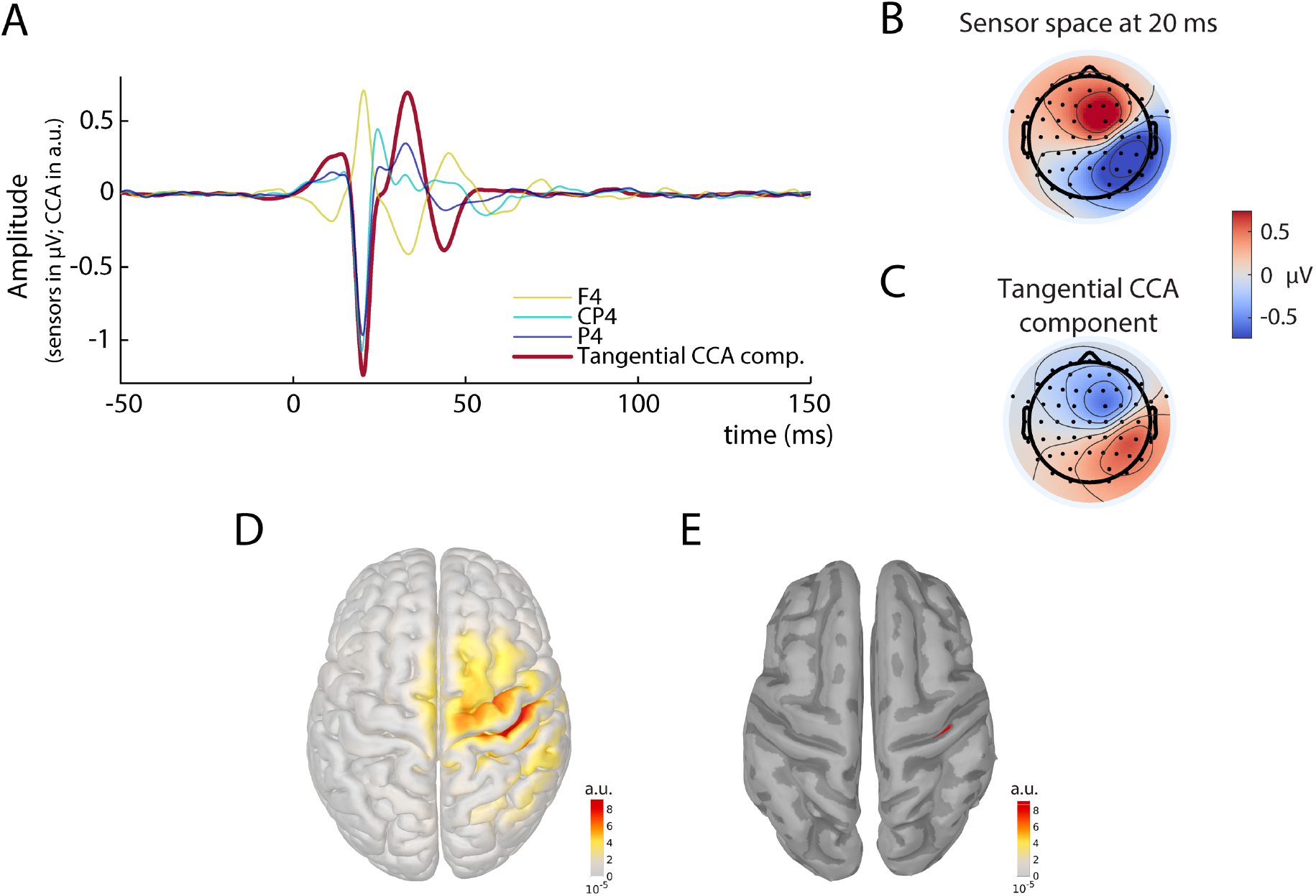
Grand average of the somatosensory evoked potential (SEP) across all subjects in sensor, CCA, and source space. A) SEP at electrodes F4, CP4, and P4, as well as its representation in the tangential CCA component. B) Scalp topography in sensor space at 20 ms post-stimulus. C) Activation pattern of the tangential CCA component. D) Sources (absolute values) underlying the spatial patterns of the tangential CCA component, reconstructed using eLoreta based on individual head models. E) Same as D but applying an amplitude threshold of 95% in order to find the strongest generators (displayed on a smoothed cortex surface).

To extract single-trial SEPs we used a variant of Canonical Correlation Analysis (CCA) in which spatial filters were trained based on a pattern matching between average SEP and single trials (Waterstraat et al., 2015; Fedele et al., 2013). With this method, we obtained a set of spatially distinct CCA components for every individual subject. In all subjects, a prominent CCA component was identified that displayed the pattern of the typical N20 tangential dipole (Fig. 2C) and showed a clear peak at around 20 ms post-stimulus (Fig. 2A). Furthermore, subsequent source reconstruction of the spatial pattern revealed that the strongest generators were located in the anterior wall of the postcentral gyrus (Fig. 2D & 2E). We focus on this CCA component in the following analyses and refer to it as the *tangential CCA component*.x

Single-trial SEPs retrieved from the tangential CCA component are displayed in Figure 3A for an exemplary subject. It is apparent that the amplitude of the early SEP fluctuates over trials, however without a clear deterministic trend (Figure 3B).

**Figure 3.**
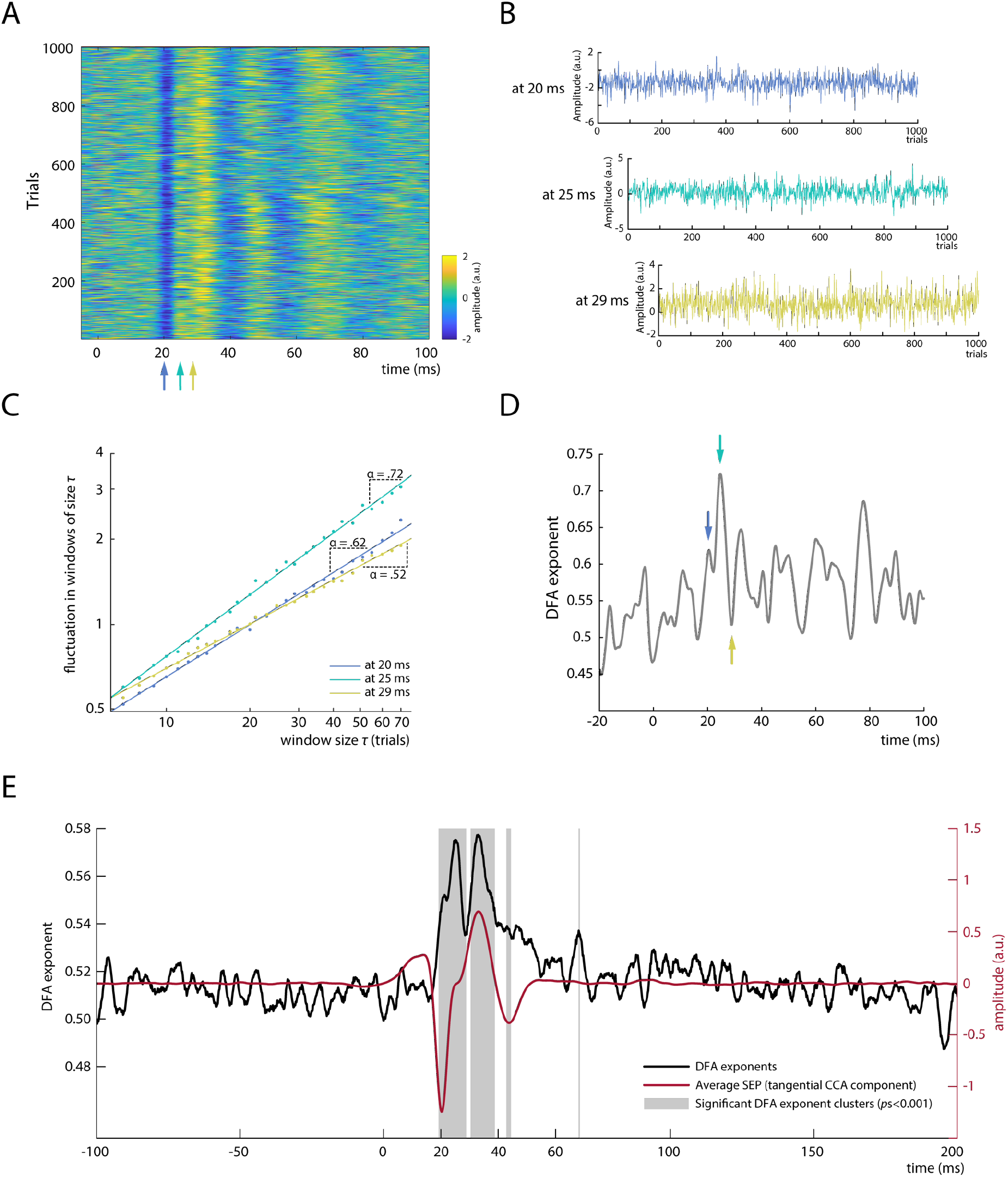
Analysis of power-law dynamics in SEP amplitude fluctuations. A) Single-trial SEPs as measured by the tangential CCA component for an exemplary subject. B) SEP amplitude fluctuations across trials for exemplary latencies (20 ms, 25 ms, and 29 ms post-stimulus). C) Detrended Fluctuation Analysis (DFA) for amplitude fluctuations depicted in B. The DFA exponent *α* is measured as the slope of a regression line fitted to the log-log relationship between window size *τ* and the fluctuation in each window size, quantifying the power-law dynamics of the signal. Note that both axes are scaled logarithmically. D) Time course of DFA exponents for an exemplary subject; blue, cyan, and yellow arrows mark the latencies which are displayed in B and C. E) DFA exponent time course averaged across all subjects (depicted in black) plotted along with the average SEP of the tangential CCA component (depicted in red). Grey boxes indicate significant clusters of DFA exponents differing from surrogate data with shuffled trial order.

### Temporal dynamics in single-trial SEP amplitude fluctuations

To evaluate the characteristics of SEP fluctuations across trials, we applied Detrended Fluctuation Analysis (DFA; Hardstone et al., 2012; Kantelhardt et al., 2001). The DFA exponent α quantifies the extent of power-law dynamics of a signal, with *α* > 0.5 indicating persistent auto-correlations; whereas *α* = 0.5 would suggest a signal without a temporal structure (i.e., white noise). DFA was performed on the amplitudes at every latency relative to the stimulus onset, across time windows of 7 to 70 trials (i.e., equivalent to around 5 to 50 seconds; exemplarily illustrated in Fig. 3B & 3C). Applying DFA at all latencies provided a *DFA exponent time course* for every subject (Fig. 3D) that indicates which portions of the early SEP show power-law dynamics (i.e., DFA exponents > 0.5). Subsequently, DFA exponent time courses were averaged across participants (Fig. 3E). The average explained variance of the power-law relationships at all latencies of the time range between 10 and 50 ms was R^2^ > .99, indicating a near-perfect fit of the DFA method for this data.

Increased DFA exponents were observed particularly in the early part of the SEP with an onset around the latency of the N20 component, whereas surrogate data generated by shuffling the trial order yielded DFA exponents close to *α* = 0.5. Two prominent peaks in the DFA exponent time course are visible from Figure 3E, with DFA exponents of *α* = .575 and α = .577, at latencies of 25 and 33 ms post-stimulus, respectively. Note, however, that the absolute value of DFA exponents highly depends on the signal-to-noise ratio (SNR) of the signal, as is further examined in simulations below which suggest DFA exponents of at least *α* = .63 when the SNR bias is taken into account. The observation of two prominent DFA exponent peaks was statistically confirmed as two main clusters were found around these two peaks by cluster-based permutation tests (*p*_scluster_ < .001). The DFA exponents were characterized by a similar yet not identical time course as compared to the magnitude of the SEP (Fig. 3E). Although the first significant DFA exponent cluster emerged together with the peak of the N20 component, the first DFA exponent peak appeared slightly later. This suggests that long-range temporal dependencies were not most pronounced at the N20 peak but rather while the potential returned back to baseline. Yet, this is not contradicting the notion of power-law dynamics in fluctuations of cortical excitability since recent evidence from pharmacological studies suggests that excitatory processes of the N20 component dominate even until the rising flank of the P35, the next component after the N20 in the SEP (Bruyns-Haylett et al., 2017).

The second DFA time course peak co-occurred with the second prominent peak of the SEP, the P35 component. This second DFA peak most likely reflects activity propagated from the N20 component to the P35 as these two components moderately covaried in our data, *β*_fixed_ = −.378, *t*(29.800) = −9.342, *p* < .001, as tested by a random-slope linear-mixed-effects model with *P35 peak amplitude* as dependent variable, *N20 peak amplitude* as independent variable and *subject* as random factor. Similarly, the two smaller clusters of increased DFA exponents at around 44 and 68 ms (Fig. 3E) may reflect the propagation of dynamics in earlier SEP components to later processing stages.

Temporal filtering in the preprocessing of the data cannot have caused these long-range temporal dependencies since (1) no DFA exponent increases were observed during the pre-stimulus baseline of the SEP and (2) additional control analyses did not show increased DFA exponents when applying the same preprocessing to stochastically independent SEP fluctuations (as tested with simulated data).

### Do power-law dynamics originate from the neuronal fluctuations in the periphery or at the thalamic level?

To investigate whether the observed temporal dynamics in cortical SEPs may arise from fluctuations in peripheral nerve excitability, we applied the same procedure as described above to the compound nerve action potential (CNAP) of the median nerve measured at the inner side of the upper arm. As expected, the nerve potential peaked at around 6 ms post-stimulus and fluctuated over trials (Fig. 4A). However, no increased DFA exponents were observed (Fig. 4B).

**Figure 4.**
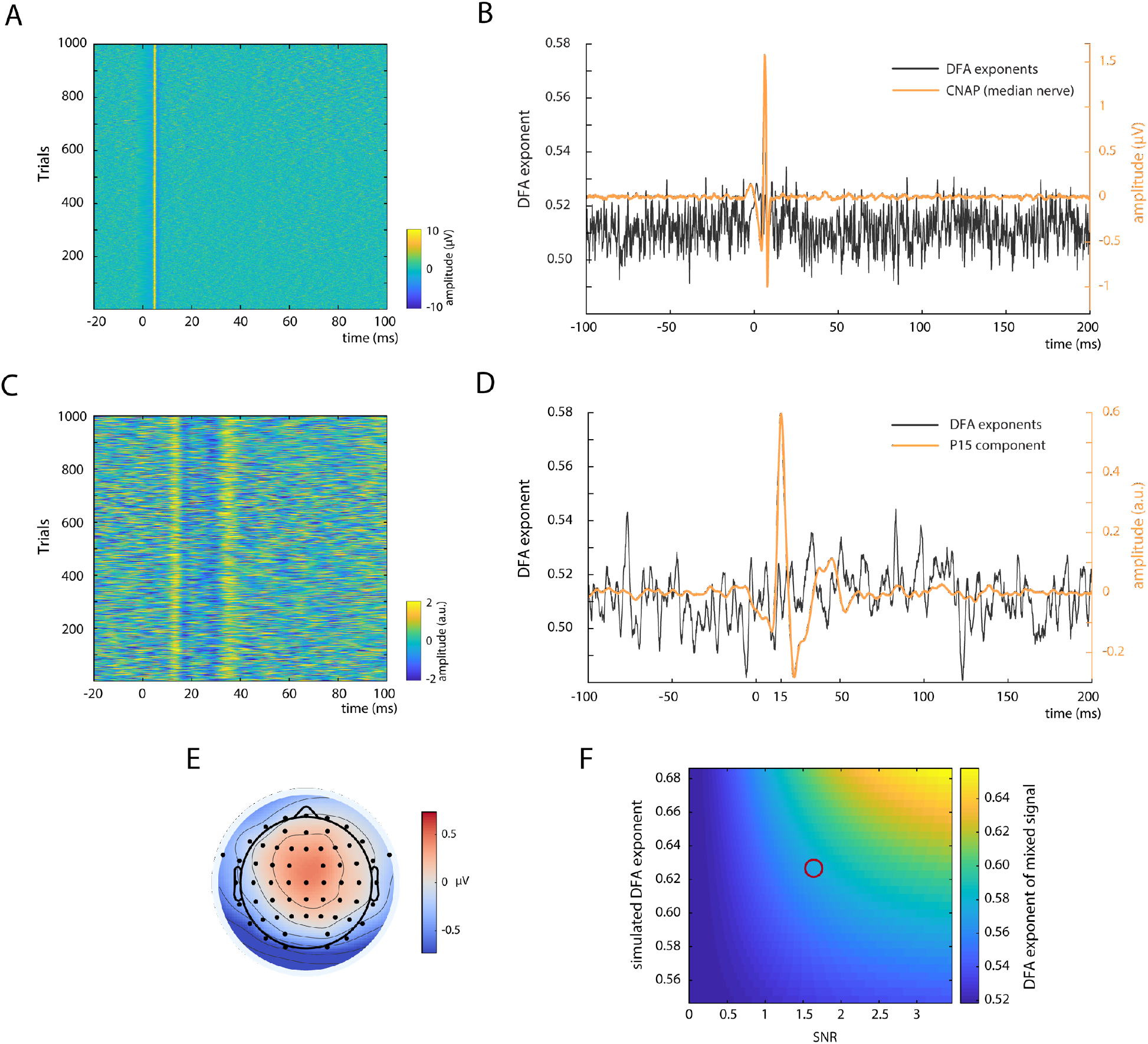
Control measures. A) Single-trial compound nerve action potentials (CNAP) of the median nerve measured on the inner side of the upper arm; depicted for an exemplary subject. B) DFA exponents (black) and CNAP (orange) of the median nerve; averaged over all subjects. C) Single-trial SEPs of the thalamus-related CCA component of an exemplary subject. D) DFA exponents (black) and SEP (orange) of the thalamus-related CCA component; averaged over the 13 subjects in which a peak at around 15 ms was observed on single-trial level. E) Average spatial pattern of the thalamus-related CCA components; averaged over 13 subjects. F) Simulation of the influence of signal-to-noise ratio (SNR) on the measurement of DFA exponents. Signals with varying DFA exponents (plotted on vertical axis) were mixed with white noise (i.e., DFA exponents of ~0.5) with varying SNR (plotted on horizontal axis). The resulting DFA exponents of the mixed signals are color-coded. The red circle indicates the region of empirically observed DFA exponents of ~0.57 at an SNR of ~1.68 suggesting an underlying DFA exponent of the unmixed source of ~ 0.63. For visualization purposes, the results of the simulation are displayed only for a sub-range of DFA exponents and SNRs here.

In addition, a CCA component was identified in 13 out of the 31 participants that contained SEP activity already at 15 ms (Fig. 4C & 4D), most likely reflecting the P15 component of the SEP which is thought to represent thalamus-related activity (Albe-Fessard et al., 1986). Also, the spatial pattern of this CCA component suggested a deeper and more medial source than the tangential CCA component (Fig. 4E). Importantly, the DFA exponents of this subcortical activity did not show any increase in the range of the P15 component (Fig. 4D) thus being in contrast with the DFA exponent increase for early cortical potentials.

### DFA exponents and SNR

Since it is known from previous studies that the signal-to-noise ratio (SNR) highly affects the measurement of power-law dynamics (Blythe et al., 2014), we investigated the relationship between DFA exponents and SNR in single-trial SEPs. On average across all participants, the SNR of the tangential CCA component was 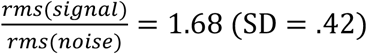 and showed a positive rank correlation with DFA exponent increase in the time range from 10 to 50 ms post-stimulus, *r* = .36, *p* = .049.

Additionally, we further clarified this relationship with simulations: We mixed signals expressing different DFA exponents with white noise (DFA exponent *α* = 0.5), for a range of SNRs, and measured the DFA exponent of these mixed signals. As is visible from Figure 4F, DFA exponents of the mixed signals are attenuated towards *α* = 0.5 when lowering the SNR. Given an SNR of 1.68 and an empirical DFA exponent of *α* = 0.575, as was the case for the tangential CCA component in the present study, our simulations suggest an underlying source with a DFA exponent of *α* ≈ 0.63. Yet this value most likely still underestimates the “true” power-law dynamics of the system as the signal term contained in the empirical estimate of the SNR is not noise free but a mixture of both signal and noise. This leads to an overestimation of the SNR and in turn to an underestimation of the degrading impact of noise on the scaling exponent.

To relate this simulation also to the others measures for which we calculated DFA exponents, we calculated the SNR of the CNAP at the upper arm and thalamic CCA components. Here, we found SNRs of 2.20 (SD = .85) and 1.33 (SD = .11), respectively, suggesting that our signal quality was sufficient to detect DFA exponent increases if they had been there since the SNR of the CNAP was even higher than that of the SEP and the SNR of the thalamic CCA component was just slightly lower.

### Power-law dynamics in alpha band activity and its relation to the SEP

Since previous studies on cortical excitability in M/EEG focused on oscillatory activity in the alpha band, we investigated both its correspondence to the early part of the SEP as well as its DFA exponents.

To test the relationship between alpha oscillatory activity and SEP amplitude, we performed a regression analysis between the mean alpha amplitude in a pre-stimulus window from −200 to −10 ms and the peak amplitude of the N20 component. Alpha activity was extracted from the same neuronal sources as the SEP by applying the spatial filter of the tangential CCA component. A significant negative relationship was found on group-level using a random-slope linear-mixed-effects model, *β*_fixed_ = −.034, *t*(25.095) = −4.895, *p* < .001. Thus, higher pre-stimulus alpha activity was associated with more negative N20 peak amplitudes (Fig. 5A).

**Figure 5.**
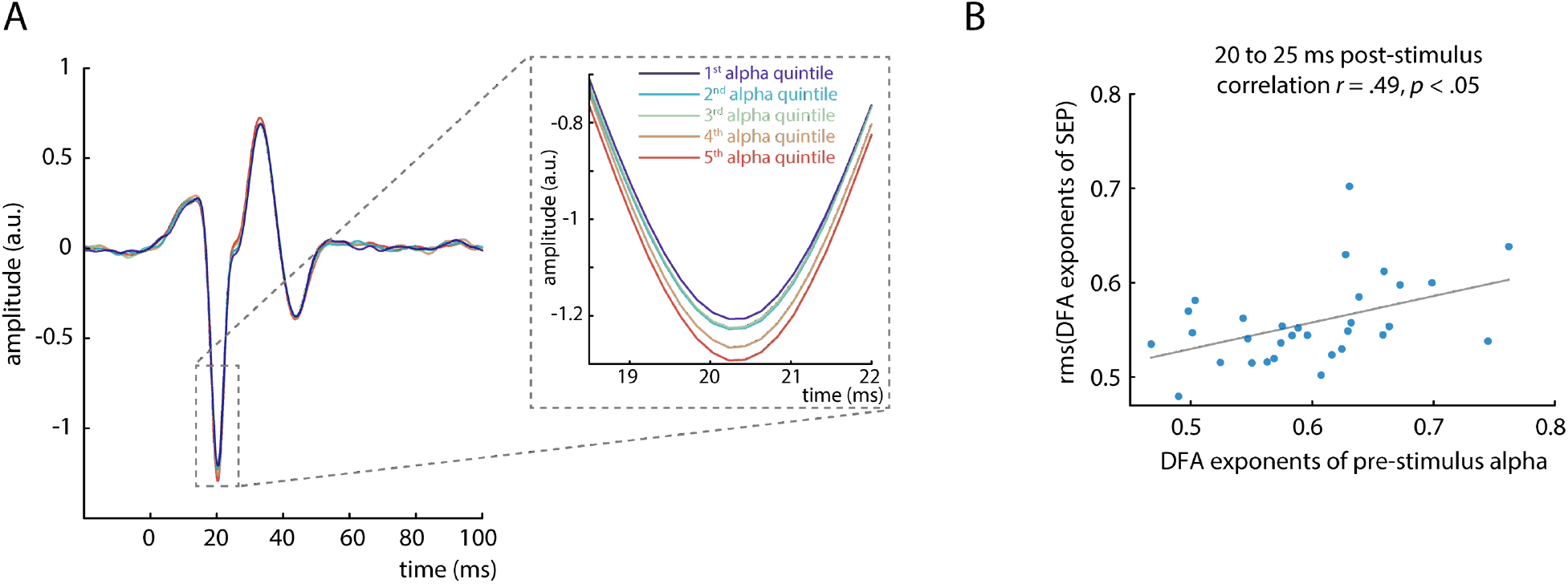
Relation between pre-stimulus alpha band activity and the early SEP. A) Average SEP (tangential CCA component) plotted by quintiles of pre-stimulus alpha band amplitude, demonstrating their relationship with the N20 component peak (inlay). B) Correlation between DFA exponents of pre-stimulus alpha amplitude and DFA exponents of the SEP in the time window from 20 to 25. The DFA exponents of the SEP were aggregated over time points using root-mean-square (rms).

To control for spurious covariation caused by the auto-correlated structure of both signals, we additionally ran permutation tests using surrogate data with comparable temporal structure as suggested in (2015). Aggregated on group-level, these tests confirmed the negative relationship between pre-stimulus alpha activity and N20 peak amplitude, *r*_group-level_ = −.035, *p* < .001.

Next, we investigated the DFA exponents of mean pre-stimulus alpha amplitude across trials. Averaged across subjects, we observed a mean DFA exponent of *α* = .60, which significantly differed from DFA exponents for shuffled trial order, *t*(30) = 6.627, *p* < .001. Also, DFA exponents in continuous, ongoing alpha activity were significantly increased across subjects, *α* = .66, *t*(30) = 10.591, *p* < .001. Thus, power-law dynamics were present in both prestimulus and continuous alpha activity.

To further test the relationship between pre-stimulus and SEP dynamics, we correlated DFA exponents of pre-stimulus alpha amplitude and DFA exponents of the SEP across participants. DFA exponents of the SEP were aggregated (using root-mean-square) in four consecutive time windows of 5 ms each, between 20 and 40 ms post-stimulus. DFA exponents of alpha activity were correlated with the DFA exponents of the first time window from 20 to 25 ms, *r* = .485, *p* = .025 (Bonferroni-corrected; Fig. 5B). However, this relationship did not emerge for any other time window between 25 and 40 ms, *p*s > .3. Notably, the SNR of the SEP cannot explain the relation between DFA exponents of alpha activity and DFA exponents of the SEP, as no relationship was found between SNR of the SEP and DFA exponents of prestimulus alpha activity, *r* = .208, *p* = .261.

Taken together, both amplitude and temporal structure of oscillatory activity in the alpha band thus relate to the corresponding characteristics of the early SEP responses, establishing a link between these two measures of instantaneous cortical excitability.

## 4 Discussion

In the present study, we investigated the temporal dynamics of neuronal excitability in the human primary somatosensory cortex by short-latency somatosensory evoked potentials. Fluctuations of excitability demonstrated power-law dynamics across trials, extending the previous notion that neuronal systems operate close to a critical state (Beggs and Plenz, 2003; Poil et al., 2012; Priesemann et al., 2013; Linkenkaer-Hansen et al., 2001; Palva et al., 2013) to variability in stimulus-evoked responses in the human sensory system. In addition, fluctuations in pre-stimulus alpha band activity and initial cortical excitation were related through their amplitudes as well as their temporal structure. For the first time, these findings thus link critical dynamics in ongoing and evoked activity as measured non-invasively in the human EEG, and directly associate the observed power-law dynamics with variability in cortical excitability.

### What do temporal dynamics in SEPs tell about the functioning of the neural system?

Eearly SEP amplitudes demonstrated power-law dynamics persisting for time windows up to 50 seconds. Given that the SEP in the time range of the N20 component only reflects initial excitatory cortical processes (Bruyns-Haylett et al., 2017; Wikström et al., 1996; Nicholson Peterson et al., 1995), instantaneous excitability thus does not seem to vary stochastically independently over time (i.e. like white noise) but is characterized by long-range temporal dependencies. This means, cortical excitability at a given moment is related to its fluctuation history and contains information about subsequent dynamics.

Such dynamics have often been interpreted within the hypothesis that the underlying system is poised at a critical state, that is a phase transition between distinct system regimes such as order and disorder (Kitzbichler et al., 2009; Sethna et al., 2001; Bak et al., 1987; Bak et al., 1988; Beggs and Plenz, 2003; visualized using the Ising model in Figure 1A). As the dynamic range, information processing and memory capacity of a system are maximized at such a phase transition (Kinouchi and Copelli, 2006; Shew and Plenz, 2013), it may be beneficial for neural systems to be tuned to this close-to-critical state although the variability inherent to it may impair an exact mapping of stimulus features and neuronal activity in sensory processing.

Typically, the power-law relationships in models of complex systems as well as in empirical neuronal avalanche recordings have been measured in the spatial domain, such as the distribution of size and duration of neuronal avalanches. Yet, critical systems should also express power-law dynamics in the temporal domain as shown for the Ising model (Zhao et al., 2017). Thus, the observed long-range *temporal* dependencies in cortical excitability in our data might in fact correspond to near-critical dynamics in the *spatial* domain as observed by Beggs and Plenz (2003) in their seminal study on the scale-free behavior of spontaneous neuronal avalanches in slices of the rat somatosensory cortex.

Specifically, the temporal power-law dynamics in our data could reflect that neuronal excitability spatially differs across the network, in agreement with the notion of scale-free neuronal avalanches, which in turn leads to the observed amplitude fluctuations in the early SEP over time. Whereas a recent approach to measure neuronal avalanches from thresholded broad-band EEG data supported the notion of critical dynamics also after stimulus presentation (Arviv et al., 2015), our findings of power-law dynamics in early SEPs demonstrate near-critical dynamics for the first time in primarily stimulus-related processes in the human brain and suggest fluctuations of cortical excitability to be the driving underlying mechanism.

### Dissociation of temporal dynamics in the cortex from peripheral and subcortical variability

To corroborate the notion of the observed power-law dynamics being a cortical phenomenon, we examined their origin in more detail.

First, we measured power-law dynamics at the onset of the N20 component (not earlier), giving no cause to assume generators of power-law dynamics in stimulus processing up-stream of the primary somatosensory cortex. Second, source reconstruction confirmed that the strongest generators of the N20 lay in the anterior wall of the post-central gyrus (Fig. 2E) as is expected from the literature for the N20 component (Allison et al., 1991). Third, no power-law dynamics were present in peripheral variability as measured from the compound nerve action potential (CNAP) of the median nerve. Fourth, thalamus-related activity reflected in the P15 component of the SEP in a subsample of 13 subjects did not show any power-law dynamics either, suggesting that neuronal variability is stochastically independent even at final subcortical processing stages.

Hence, we conclude that the observed power-law dynamics most likely are of cortical origin.

### Underestimation of power-law dynamics due to signal-to-noise ratio

It is known that the signal-to-noise ratio (SNR) has an impact on the estimation of DFA exponents (Blythe et al., 2014), that is, even a signal with an exponent of *α* ≈ 1.0 would result in lower DFA exponents when being contaminated with strong noise characterized by exponents of *α* ≈ 0.5. Given our empirical single-trial SNR, simulated scenarios with varying SNRs and DFA exponents suggested a lower bound for the underlying DFA exponent of *α* ≈ 0.63, which is in the range of temporal power-law dynamics reported in previous MEG/EEG studies for ongoing alpha band activity (Linkenkaer-Hansen et al., 2001; Palva et al., 2013).

### Relationship between pre-stimulus alpha activity and initial cortex excitation

Following the idea that oscillatory activity in the alpha band reflects cortical excitability (Romei et al., 2008; Klimesch et al., 2007; Zrenner et al., 2017; Sauseng et al., 2009), we tested whether this measure in a pre-stimulus window was related to the initial cortex excitation as assessed by the N20 amplitude. In our data, lower amplitudes of alpha band activity, hypothesized to reflect a state of increased excitability, were associated with smaller (less negative) N20 amplitudes. Although at a first glance this finding seems to contradict the hypothesis of low alpha activity being associated with higher excitability (Jensen and Mazaheri, 2010; Klimesch et al., 2007) it may be explained in a straightforward manner by the neurophysiological basis of EEG generation.

The scalp EEG reflects relative changes in collective charge distributions resulting from neuronal activation manifested in primary post-synaptic currents (PSCs; Lopes da Silva, 2004; Kandel et al., 2000; Ilmoniemi and Sarvas, 2019). The magnitude of an EEG potential *U* emerging through synchronous activity of a well-specified neuron population, as is assumed for the N20 component of the SEP, should follow the general relationship

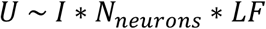

where *I* denotes the sum of local primary post-synaptic currents, *N_neurons_* the number of involved neurons, and *LF* the lead field coefficient projecting source activity to the electrodes on the scalp. Since *N_neurons_* and *LF* can be assumed to be constant for the N20 component in a sequence of unchanging median nerve stimuli, we believe that primarily *I*, reflecting excitatory PSCs, contributed to the amplitude variability of the early part of the SEP.

Now, assuming that states of higher neuronal excitability are associated with membrane depolarization on a cellular level, the electrical driving force for further depolarizing inward trans-membrane currents is decreased and less current is needed to reach the threshold potential for excitatory responses (Castro-Alamancos, 2009). This leads to *decreased* PSCs at high excitability states (Deisz et al., 1991) and would result in lower amplitudes in the EEG. Hence, one should rather expect *decreased* N20 components following *low* pre-stimulus alpha activity, as was the case in our data.

The relationship between pre-stimulus alpha activity and early SEP amplitudes is further corroborated by their corresponding degree of power-law dynamics in the time range of the SEP from 20 to 25 ms post-stimulus (Fig. 5B). Intriguingly, the idea that this link is established via cortical excitability manifested in pre-stimulus membrane potentials is consistent with a recent study which indeed demonstrated near-critical dynamics in membrane potential fluctuations in the turtle visual cortex (Johnson et al., 2019).

Although this notion should be treated with some caution as it is based on the strong assumption that mechanisms observed in single-cell recordings (in animals) can be generalized to cell populations in the human cortex, the present findings could represent the missing link between power-law dynamics on micro (single-cell) and macro (cell population) scale and would relate findings of criticality in neuronal avalanches to non-invasively measured EEG potentials in humans.

### Implications for the perspective on neural variability

Why neuronal systems express large variability, particularly in perceptual processes, has been an enduring question for many years. The perspective on the temporal structure of variability in neural responses now extends our understanding of the underlying organizing principles. Not only is stimulus-evoked activity dependent on pre-stimulus network states but even the network states themselves seem to fluctuate in a structured manner – in our data manifested in near-critical dynamics in instantaneous cortical excitability. These dynamics are poised just at the border between deterministic and indeterministic behavior, which may enable neural networks to adaptively adjust during stimulus processing. Criticality may thus represent a compellingly parsimonious explanation of moment-to-moment fluctuations in neural responses and why they can actually be beneficial for neural systems.

## Author contributions

T.S. and V.V.N. conceptualized the study with essential input from G.W., G.C. and A.V.. T.S. collected and analyzed the data in coordination with V.V.N.. S.H. and G.W. contributed central analysis tools. T.S. and V.V.N. wrote the first version of the manuscript. T.S., G.W., S.H., G.C., A.V. and V.V.N. edited the manuscript.

## Acknowledgements

We thank Sylvia Stasch for participant recruitment and help with data collection, as well as Alice Hodapp for supporting the source reconstruction of the EEG.

